# SLEDGe: Inference of ancient whole genome duplications using machine learning

**DOI:** 10.1101/2024.01.17.574559

**Authors:** Brittany L. Sutherland, George P. Tiley, Zheng Li, Michael TW McKibben, Michael S. Barker

**Affiliations:** Department of Ecology & Evolutionary Biology, University of Arizona, Tucson, AZ 85705; Department of Biology, George Mason University, Fairfax, VA 22030; Royal Botanic Gardens, Kew, Richmond TW9 3AB, UK

## Abstract

Ancient whole-genome duplication--previous genome duplication events that have since been eroded via diploidization, are increasingly identified throughout eukaryotes. One of the constraints against large-scale studies of ancient eukaryotic WGD is the relatively large, high-quality datasets often needed to definitively establish ancient WGD events; alternatively, the more low-input method interpretation of genome-wide synonymous substitution rates (Ks plots) is prone to bias and inconsistency. We improve upon the shortcomings of the current Ks plot method by building a Ks plot simulator. This data-agnostic approach simulates common distributions found in Ks plots in the presence or absence of ancient WGD signatures. In conjunction with a machine-learning classifier, this approach can quickly assess the likelihood that transcriptomic and genomic data bear WGD signatures. On independently-generated synthetic data and real plant transcriptomic data, SLEDGE is capable of correctly identifying ancient WGD in 93-100% of samples. This approach can serve as a quick classification step in large-scale genomic analyses, identifying putative ancient polyploids for further study.

## Introduction

Polyploidy, or whole genome duplication (WGD), has been an important factor throughout eukaryote evolution. Extant polyploidy is widespread among diverse eukaryotic lineages. Approximately 30% of plants are recent polyploid species (Barker *et al*., 2016; Wood *et al*., 2009; Mayrose *et al*., 2011), and polyploidy is common in some animal lineages, such as weevils (Stenberg *et al*., 2003), stick insects (Myers *et al*., 2013), parthenogenetic reptiles (Sexton, 1980), and salmonid fishes (Comber *et al*., 2004). Paleopolyploidy, ancient WGDs that may not be readily discernible in extant lineages, is known to have influenced the evolution of most eukaryotes. Nearly 250 ancient WGDs have been identified in plants (One Thousand Plant Transcriptomes Initiative [OTPTI] 2019; Li and Barker 2020; Ren et al. 2018; Wendel 2015; Barker et al. 2016; Smith et al. 2018), fungi (Albertin and Marullo 2012; Campbell et al. 2016; Wolfe and Shields 1997), and animals (Li et al. 2018; Berthelot et al. 2014; Kenny et al. 2017; Shingate et al. 2020; Liu et al. 2021; Farhat et al. 2023; Conant 2020; Gundappa et al. 2022; Redmond et al. 2023) including at least two rounds of WGD that occurred during the evolution of vertebrates (Ohno 1970; Panopoulou and Poustka 2005; Blomme et al. 2006).

Ancient WGDs are commonly inferred by identifying peaks of gene duplication from age distributions of paralogs within a single species (Blanc and Wolfe 2004; Cui et al. 2006; Schlueter et al. 2004; Tiley et al. 2018; Barker et al. 2008; Vanneste et al. 2013; Initiative and One Thousand Plant Tra…). The synonymous divergence (Ks) of all gene duplications in a genome is often visualized as a histogram commonly called a Ks plot. Ongoing gene birth and death from small scale duplication and loss events produce an exponential background distribution in the genomes of most species, with relatively high gene birth and death producing a steep slope (Lynch and Conery 2000). WGDs, by contrast, create a “burst” of duplicated genes. These paralogs are often retained for longer periods of time than small scale duplications, due to the length of time required to remove the large number of duplicates through stochastic processes and because dosage balance retains many paralogs to maintain the stoichiometry of their gene products (Freeling 2009; Tiley et al. 2018; Conant et al. 2014; Mayfield-Jones et al. 2013; Vanneste et al. 2013). The divergences of paralogs from a WGD yield a localized “peak” in the Ks plot that corresponds to the approximate age of the duplication event. Previous analyses have found that inferences of WGDs from Ks plots are consistent with synteny analyses from sequenced genomes (Li and Barker 2020), but with the advantage of presenting a lower barrier to analysis because only assembled genes are needed rather than assembled chromosomes or large contigs. For this reason, Ks plots can be built from either genomic or transcriptomic data. Despite these positives, Ks plot analysis is often more art than science. Although a variety of methods have been developed to identify significant peaks in Ks plots (Schlueter *et al*., 2004; Cui *et al*., 2006; Barker *et al*., 2008), none of these approaches explicitly model a WGD. Ks plots from older WGD events, from taxa with lower than usual rates of duplicate gene retention, or generated from poorer quality transcriptomic data can present ambiguous peaks, which may be interpreted differently by different practitioners. Efforts to make the analysis of Ks plots more objective have met with limited success. The current state of the art is to fit *n* distributions in a Ks plot using mixture-models. Although this method accurately identifies WGD-related peaks in the distribution, it also overestimates peak number, calling spurious peaks that do not correspond to WGDs (Tiley *et al*., 2018; Vanneste *et al*., 2013).

The advent of machine learning presents a new avenue for the inference of WGDs from Ks plots. Binary classification of features from noisy, variable, or incomplete data is a problem for which machine learning is uniquely suited. By developing a dataset that represents WGDs under various conditions, machine learning algorithms can be trained to detect the presence or absence of WGD across a wide range of taxa and WGD ages. We developed new tools for rapidly simulating gene age distributions with and without WGDs, trained models with four different machine learning algorithms, and tested performance of these models against a large simulated dataset and two empirical datasets. Here, we release the best-performing models and simulation code as the Supervised Learning Estimation of Duplicated Genomes (SLEDGe) package, which accepts gene age distributions generated from tools such as DupPipe (Barker *et al*., 2010) or wgd (Zwaenepoel and Van de Peer 2018) to quickly and accurately classify genomes with and without putative ancient WGDs across a range of taxa, WGD ages, and data qualities.

## Methods

### Generating simulated gene age distributions

To train machine learning models to detect ancient WGDs, we simulated gene trees within a single individual under a simplified duplication and loss procedure. Simulations were designed to mirror Ks distributions observed in empirical data, but were otherwise agnostic to underlying biological processes (Fig. 1). The simulation runs forward in time, where the branch length measured in Ks of each gene tree root is drawn from a gamma distribution. Genes are duplicated at rate µ_1_ and lost at rate λ_1_. Gene trees can be multicopy at the root, which is drawn from a geometric distribution. Duplication and loss occurs in discretized Ks intervals of 0.1. The simulations make the assumption that a Ks of 0.001 approximates one million years of evolution (e.g. Blanc and Wolfe 2004; Gaut et al. 1996). µ and λ are the probability of a duplication or loss event along a branch per million years (e.g. Zwaenepoel and Van de Peer 2019; Tiley et al. 2016; Hahn et al. 2005). The duplication and loss process is binned into 0.1 intervals to speed simulations where many replicates were needed. For simulating a WGD, the time before present in Ks can be specified where a fraction of lineages, *q*, are duplicated. To generate the characteristically sharp peaks near Ks of 0 observed in many empirical studies, we allow for a shift in duplication and loss rates, µ_2_ and λ_2_, at a time determined *a priori* where an investigator can, for example, decrease loss rates towards the present. Such behavior is consistent with empirical observations from plant genomes (Rody et al. 2017; Li et al. 2016) and motivated by recent analyses where simulating codon evolution on gene trees with constant duplication and loss rates among branches had difficulty capturing features of empirical Ks plots (Tiley et al. 2018). We incorporated rate variation into gene trees by drawing individual branch rates from a gamma distribution. In total, a master dataset of approximately 200,000 simulations (100,000 with WGD [“positives”] and 100,000 without WGD [negatives”]) was generated.

**Fig 1:**
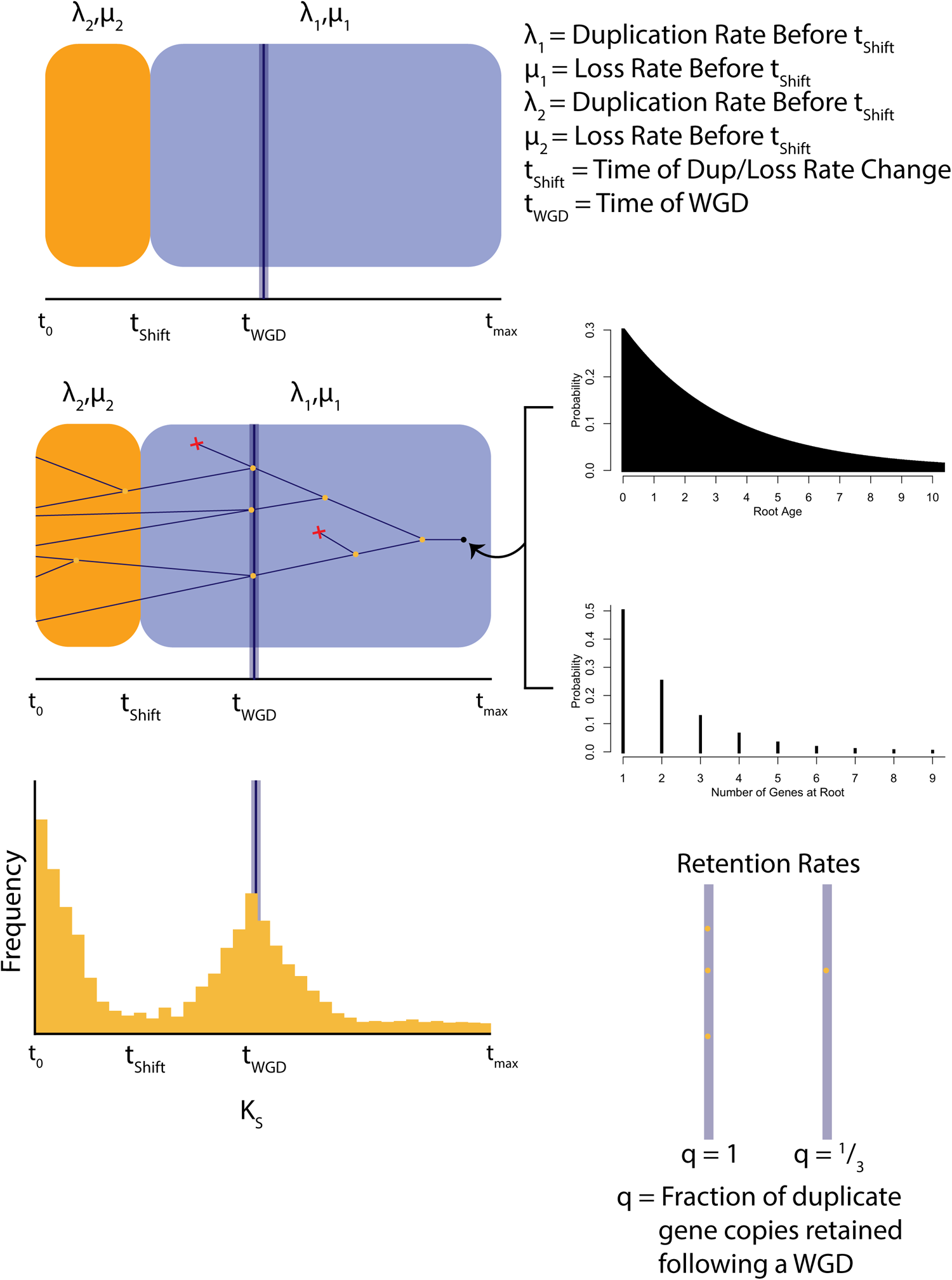
Diagram of Ks data simulation method. a) Two sets of retention and loss rates were modelled to allow for sharper Ks peaks near Ks=0 (orange) and wider peaks at longer times before present (blue). b) The number of genes at the root of each tree was generated from a gamma distribution, and the number of duplicates created at each time point was determined by µ_1,2_ and λ _1,2_. c) Synonymous substitution rates across all retained duplicates within all gene tree was plotted to generate each Ks plot. d) The fraction of all genes within a genome duplicated at time t is modeled by q.

With the same tools used to generate our training datasets, a test dataset of simulated Ks distributions was independently generated. All permutations of simulation conditions (birth/death rates and Ks values) were generated once each for positive (*q*=1) and negative (*q*=0) simulations, creating a test dataset of 3,204 simulations.

### Training models

Because training runtime increased exponentially with training dataset size, the master dataset was subsampled at 20%, 40%, 60%, and 80% density to generate alternative classification models. Preliminary analyses suggested that height and variance of two critical features of each distribution—the background duplication curve and the WGD peak—varied substantially with histogram resolution. To assess the effect of resolution on model performance, we chose a range of histogram resolutions from 99 to 40,099. For most algorithms, histogram bin number was tested every 100 bins between 99 and 20,099 bins, and every 1,000 bins thereafter.

Testing and training of classification models was done using the scikit-learn Python package (Pedregosa, 2011) and implementing the Support Vector Machines classifier with a Radial Basis Function kernel. The algorithm was trained for every selected bin number for the 20k, 40k, and 60k training dataset sizes and up to 20,099 bins for the 80K and full dataset sizes due to runtime constraints. Collectively, 1,020 models were trained.

### Evaluating trained models

We evaluated the performance of each trained model using two different test datasets, one dataset of simulated samples and one dataset of empirical, synteny-confirmed samples. Because the Ks distribution simulator does not produce Ks values of precisely zero, any values of Ks = 0.0 were removed from empirical Ks distributions prior to testing. The empirical dataset was chosen *a priori* because it represented different compositions of data quality and WGD frequency. Models were tested against 91 taxa taken from the 1000 Plant Transcriptome Project (1KP; OTPTI 2019) for which syntenic data exists for either that taxon or a congener. Each of these taxa have well-represented, high quality transcriptomes and a well-annotated genome. Because synteny analysis is considered the “gold-standard” for diagnosing ancient WGD, this dataset was considered to have the best supported examples of WGD peaks in Ks plots and was the most valid test of model performance.

In addition to the simulated and empirical syntenic datasets, we also compared SLEDGe classifications to predictions made via other means on two other empirical datasets. First, we tested the congruence of SLEDGe classifications with other methods on the remainder of the 1KP dataset. Because plant datasets are unavoidably biased toward having ancient WGDs (1195 out of 1395 in all of 1KP and 82 of 91 in the syntenic subset contain ancient WGD events), we also tested against a combined plant and insect dataset that reflected a more balanced composition of positive and negative samples– 47.6% with WGD, 52.4% without.

### Evaluating the recall of SLEDGe models to partial genome duplication

To assess the likelihood that SLEDGe models may mis-classify large segmental duplications or aneuploidy events as WGDs, an additional simulated test dataset was developed. These simulations differed from existing positive simulations in that, rather than simulating gene duplications in 100% of gene families at the time of WGD, a random subset of gene families contained duplications. Subsets ranged from 100% of gene families (equivalent to previous positive simulations) to 1% of gene families, and were simulated at 10% intervals (10% - 100%) plus two additional datasets at 5% and 1%. Simulated datasets were analyzed using SLEDGe, and the proportion of samples classified as positive for WGD was recorded.

### Implementation details and description of software input/output

SLEDGe is free and open source software implemented in Python 3.3 using the scikit-learn machine learning package. SLEDGe requires two to four input arguments. SLEDGe requires one input directory of sample Ks values. Each sample file may be formatted as either one column containing all Ks values or as output of the DupPipe pipeline. To save the output, a text file is specified using the -o flag. In this mode, the default Ks histogram resolution and default model are assumed and implemented. If the user wishes to change the histogram resolution or model, those may be specified using the -h and -w flags, respectively. SLEDGe outputs a text file of each filename from the input directory followed by a positive or negative classification.

## Results

### Effects of training dataset size and histogram resolution on classifier performance

Overall model performance and the effects of Ks histogram resolution and training dataset size were assessed against the simulated testing dataset, which consisted of 1,702 histograms each with and without WGDs. Performance of each algorithm was measured by calculating recall (in this context, the ability of a model to accurately classify positive WGDs), precision (ability of a model to accurately classify negative WGDs), and F_1_ score (the harmonic mean of recall and precision and a measure of overall accuracy). Each metric ranges from 0 to 1, with 1 reflecting a perfectly discriminating model. Overall, the four test algorithms varied considerably in their classification ability, and within some algorithms, training dataset size and Ks histogram resolution had a marked effect on performance.

On the simulated dataset, all algorithms except NaïveBayes had generally good performance, with F_1_ scores of 0.90-0.95. Exceptions to this were poor performances of the NaïveBayes algorithm (F_1_: 0.51-0.64; Fig. 2C) and the SVC classifier when trained on the smallest training set (F_1_: 0.45-0.52; Fig. 2A). Both SVM and SGD performed well aside from the lowest resolutions, with F_1_ scores of 0.9-1.0; however, SGD accuracy slowly decreased with higher histogram resolutions (i.e., more bins). The lowest histogram resolutions of the SVM classifier achieved perfect resolution of positive and negative WGDs (F_1_ = 1.0). Recall and precision followed similar patterns for all four algorithms, with NaïveBayes exhibiting the lowest overall performance (Recall = 0.38-0.49, Precision = 0.70-0.78; Fig. S1, S3) aside from the 20K SVC models (Recall = 0.00-0.32, Precision = 0.00-0.50; Fig. S1, S3).

**Fig 2:**
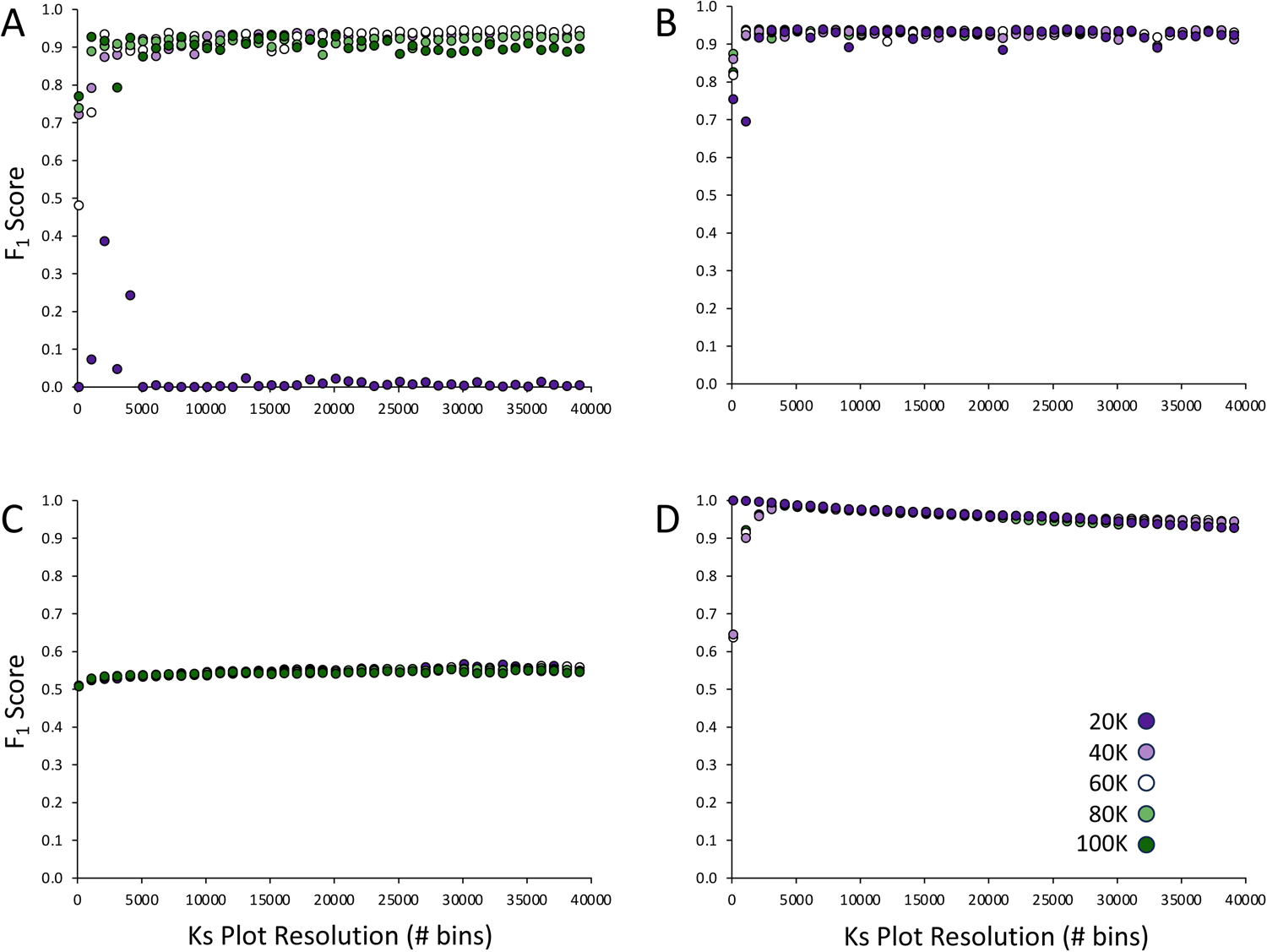
F_1_ scores of trained SLEDGe models against a test dataset comprising 3402 simulated Ks distributions (1701 displaying only background duplication rate, 1701 displaying both background duplication and one WGD). Tested models were trained used four classification algorithms and five training dataset sizes. Classification algorithms were A) LinearSVM, B) Stochastic Gradient Descent (SGD), C) Naive Bayes, and D) Support Vector Machines.

### Performance of classification algorithms against plant syntenic data

To assess if models had different responses to real as opposed to simulated data, we likewise tested all models against the 91 empirical plant Ks distributions or which synteny analysis has been performed (Fig. 3). Performance against syntenic data dropped considerably for most (but not all) algorithms, suggesting overfitting to the training data.

**Fig 3:**
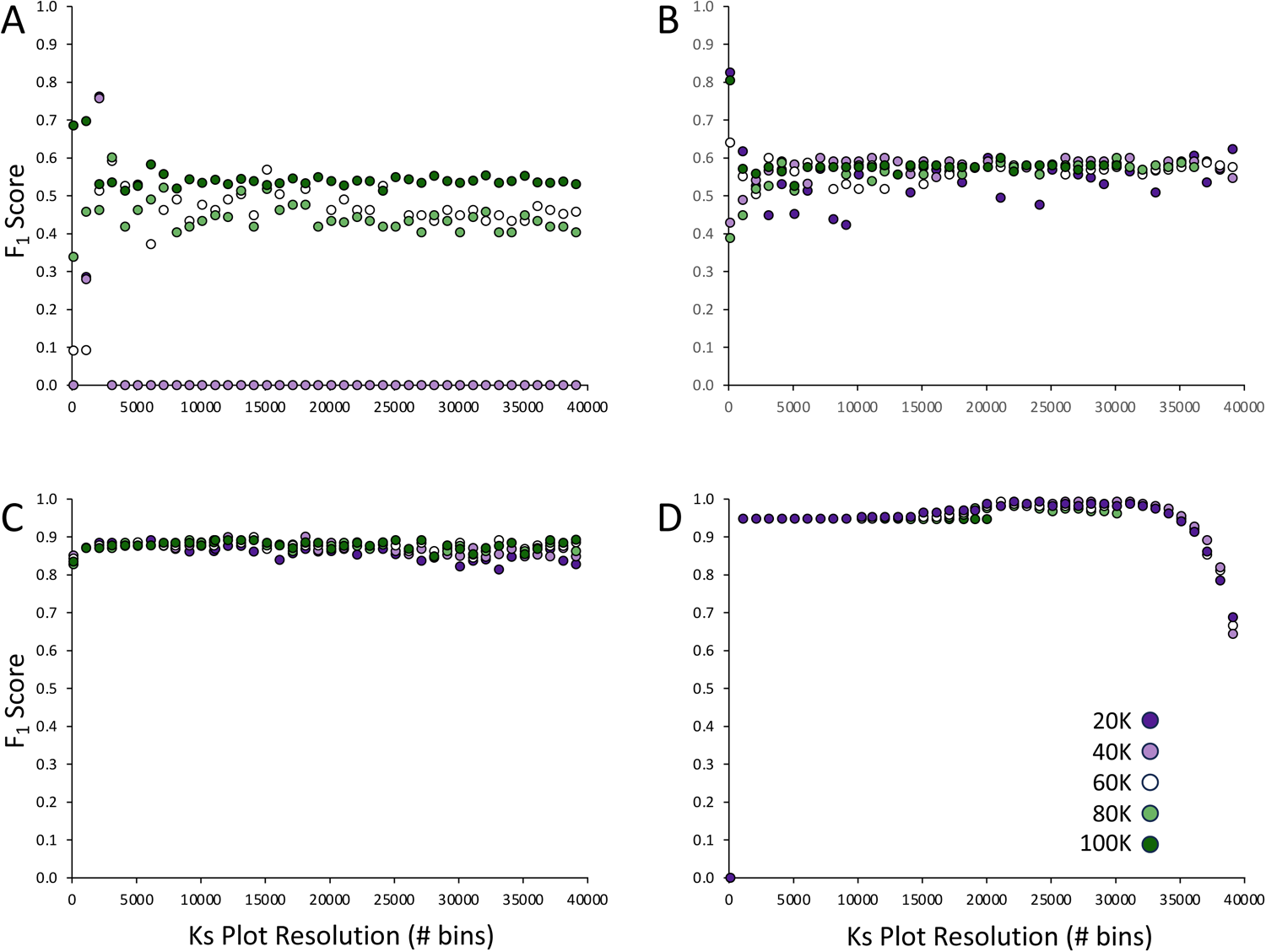
F_1_ scores of trained SLEDGe models against a test dataset comprising 91 empirical green plant Ks distributions for which synteny data confirmed classification (82 with WGD, 9 without). Tested models were trained used four classification algorithms and five training dataset sizes. Classification algorithms were A) LinearSVM, B) Stochastic Gradient Descent (SGD), C) Naive Bayes, and D) Support Vector Machines. Arrow in panel D points to model selected for SLEDGe package.

#### SVC Classifier

As with the simulated test dataset, the SVC classifier performed worst when trained on smaller training sets—it defaulted to classifying almost all samples (approx. 90% of which have known WGDs) as negative when trained on the 20K and 40K datasets. Otherwise, its average F_1_ score was 0.48. However, there was stochasticity in F_1_ score with lower bin resolutions, and accuracy reached over 50% with one set of model parameters (F_1_ score: 76% with 20K and 40K training sets and 2099-bin histogram resolution; Fig. 2).

#### SGD Classifier

The SGD classifier was only a marginal improvement over SVC on the syntenic dataset. Again, some stochasticity at low resolutions and an F_1_ score that topped out at 83% for the largest training dataset, but otherwise F_1_ for all training dataset sizes averaged 0.58. Unlike SVC, training dataset size did not have a marked effect on performance.

#### NB Classifier

The NaïveBayes classifier performed better on the syntenic dataset than it did on the simulated dataset, with an average F_1_ score of 0.88. Neither histogram resolution nor training dataset size had a large effect on performance relative to the syntenic test data. The NaïveBayes classifier also had consistently high recall and precision (Figs. S2C, S4C).

#### SVM Classifier

For the syntenic dataset, the Stochastic Vector Machines (SVM) algorithm had both the most dynamic response to Ks histogram resolution and the best overall performance. While performance appears high at low resolutions, this is an effect of the high-skewed syntenic dataset—the models merely classified all samples as positive. Likewise, at very high histogram resolutions, performance fell off dramatically. However, between 20,099 and 30,099 bins, the SVM classifier had consistently strong performance with average F_1_ values over 0.95. The classifier trained with the 40,000-sample dataset and with a resolution of 22,099 bins had an F_1_ score of 1.0, accurately classifying all samples. Because this model likewise performed well on the simulated data (F_1_: 0.96), we chose this model to proceed with further validation.

### Concordance with empirical data and effect of data quality on performance and empirical syntenic datasets

SLEDGe was further tested against a large additional empirical dataset: ∼1,400 green plant transcriptomes generated as part of the 1000 Plant Transcriptomes project (Initiative and One Thousand Plant Transcriptomes Initiative, 2019). Few samples within this dataset have specific or congeneric synteny confirmation of a WGD, but have been evaluated for presence or absence of WGDs using standard model-based approaches followed by hand-curation of Ks plots. The best-performing SLEDGe model was congruent with prior 1KP assessments 93.7% of the time.

Overall, SLEDGe was less congruent with 1KP classifications for samples in which fewer paralogs were retained, although the direction of this disagreement was not consistent. SLEDGe classification did not vary with TransRate scores (Fig S5), a measure of transcriptome quality ranging from 0 to 1 (Smith-Unna et al., 2016). However, there was no obvious pattern between retained paralog number and SLEDGe classification. The average number of retained paralogs was highest for samples scored by both SLEDGe and prior 1KP assessment as positive for WGD and lowest when both classified samples as negative (Table 1). However, the average retained paralog number in mismatches SLEDGe scored as negative was higher than the number in which both assessments agreed. This may reflect limitations of the current SLEDGe implementation, insufficient read depth in 1KP samples, or both.

**Table 1:**
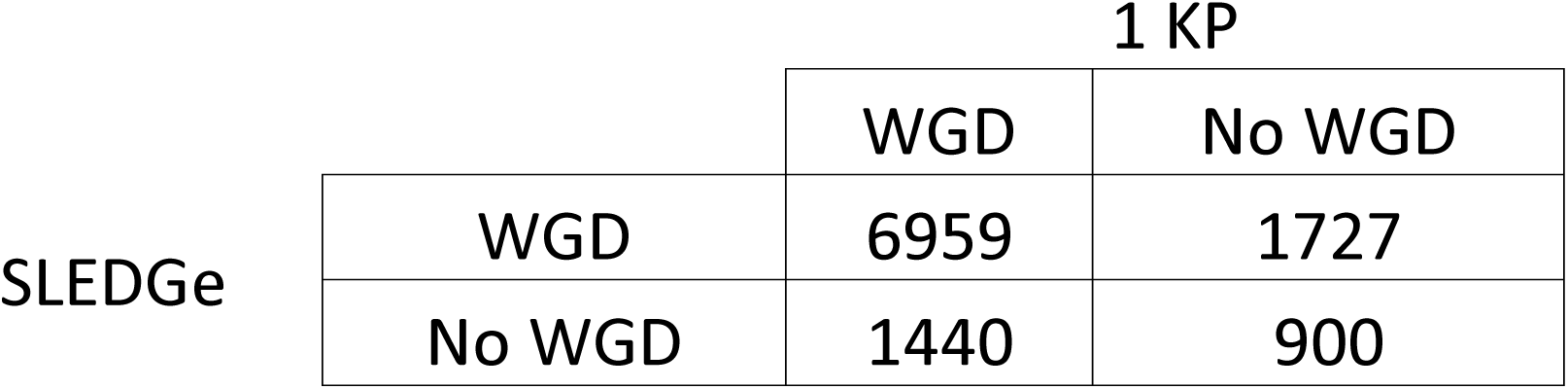
Comparison of SLEDGe and prior classifications on transcriptomes included in the 1KP dataset.

|  |  | 1 KP |  |
| --- | --- | --- | --- |
|  |  | WGD | No WGD |
| SLEDGe | WGD | 6959 | 1727 |
|  | No WGD | 1440 | 900 |

To explore the lower limit of transcriptome sample sizes for which the currently trained SLEDGe models are valid, we randomly subsampled 200 large simulated samples (>3,000 retained duplicates) to create a series of samples retaining fewer duplicate genes (250, 500, 1,000, 1,500, 2,000 duplicates, respectively). Samples with at least 1,500 duplications were correctly classified >80% of the time, whereas those with 1,000 or fewer duplications were correctly classified >60% of the time (Fig. 5).

### Effect of partial duplication on SLEDGe performance

Because partial duplications are relatively common across plants, a simulated dataset was generated to assess the performance of SLEDGe against partial genome duplications. Although most segmental duplications and aneuploidy events affect less than 10% of the genome, triploidy could, in this setting, be interpreted as a 50% genome duplication event. Our test dataset varied in initial retention from 1% to 100% gene duplicates retained, and consisted of 18,711 simulations (1,701 simulations x 11 retention levels). SLEDGe identified WGDs in over 80% of simulations for which at least 50% of the genome was duplicated, but in only 10% for which 10% or less of the genome was duplicated (Fig. 6).

## Discussion

SLEDGe provides a novel means to repeatably and rapidly infer ancient WGD events from Ks plots derived from genomic or transcriptomic data. Inferring ancient WGDs from gene age distributions, or Ks plots, has arguably been more art than science. Interpretation of peaks is confounded by biological variation that alters the signal of ancient WGD, including variation in gene birth rates, death rates, gene retention, and loss of signal as paralogs become saturated or as newer WGDs mask older events (Tiley et al. 2018; Vanneste et al. 2013). Interpretation can be further complicated by partial or segmental duplication. Statistical approaches, such as mixture models (Schlueter et al. 2004; Barker et al. 2008; Tiley et al. 2018), K-S goodness of fit tests (Cui et al. 2006), and SiZer (Chaudhuri and Marron 1999; Barker et al. 2008) are useful for identifying peaks in Ks plots, but cannot infer if such peaks in an age distribution result from WGD. SLEDGe addresses this issue by simulating ancient WGDs of multiple ages and across a range of gene birth and death rates. It provides the first model-based approach to infer WGDs in Ks plots and makes WGD interpretation more repeatable and consistent. We found that SLEDGe performed well against simulated and synteny-verified empirical data. It correctly inferred WGD events in both test datasets, and identified WGDs in concordance with another hand-curated dataset (One Thousand Plant Transcriptomes Ini…). Not only does SLEDGe provide a new tool for simulating and inferring WGDs, it also serves as proof of concept that supervised-learning approaches are a tractable approach for analyzing genome duplications in eukaryotic genomes and transcriptomes.

We assessed the utility of four algorithms commonly used for machine learning classification tasks built into the readily available machine-learning package scikit-learn. Two algorithms impose a linear discriminator to separate data into two classes, while two others were free to impose multifactorial and non-linear discriminants. Three of the four algorithms (SVC, SGD, SVM) performed well against simulated data, with mean F_1_ scores of 0.9 or better. By contrast, only one algorithm, SVM, performed well against empirical data. The ability of SVM to accurately classify both simulated and empirical data suggests that our simulator is capable of closely modeling real Ks plots.

Early work on this project suggested that histogram construction methods could substantially impact classifier performance; however, we found little difference in performance across histograms with different numbers of bins. SVM showed the greatest response to histogram resolution, with peak performance occurring between 20,099 and 30,099 bins, and a steep decline in performance at higher bin numbers. This decline is likely because there are too few data points distributed across a high number of bins, thereby making any “peak” unidentifiable. Despite the large number of bins used, peaks in positive simulations and in empirical samples containing WGD were still visible to the naked eye in most cases. We chose to use all Ks rates for a given transcriptome rather than imposing an upper-limit cutoff.

We demonstrated that the currently trained SLEDGe model has one significant limitation, likely related to the use of high bin numbers in constructing histograms. Transcriptomes or genomes that retain a relatively low number of gene duplications (∼1,000 retained duplicates) are more likely to be classified as negative for WGD. Although it is expected for samples without WGD to retain a smaller number of duplicates, we found substantial overlap between the number of duplicates retained and presence or absence of WGD, such that simply parsing by number of retained duplicates is insufficient. We urge caution when using SLEDGe on samples for which fewer than 1,000 duplicate gene families are retained from DupPipe or other methods. Training new SLEDGe models for smaller sample sizes may address this issue.

We also found that SLEDGe is sensitive to the initial proportion of the genome duplicated. Although it consistently infers WGDs in samples in which 60% or more of the genome has been duplicated, it classifies samples with 10% or less of the genome duplicated—duplication proportions similar to those found in aneuploids—as negatives (Fig. 4). For this reason, SLEDGe is unlikely to erroneously identify WGDs in samples with a history of aneuploidy but lacking other large-scale genome duplication. Notably, a triploid under our simulator counts as a genome with a 50% duplication event. Based on simulated data, the currently trained SLEDGe models appear to correctly diagnose triploids as polyploids and are much less likely to misdiagnose smaller partial duplication events as polyploidization.

**Fig 4:**
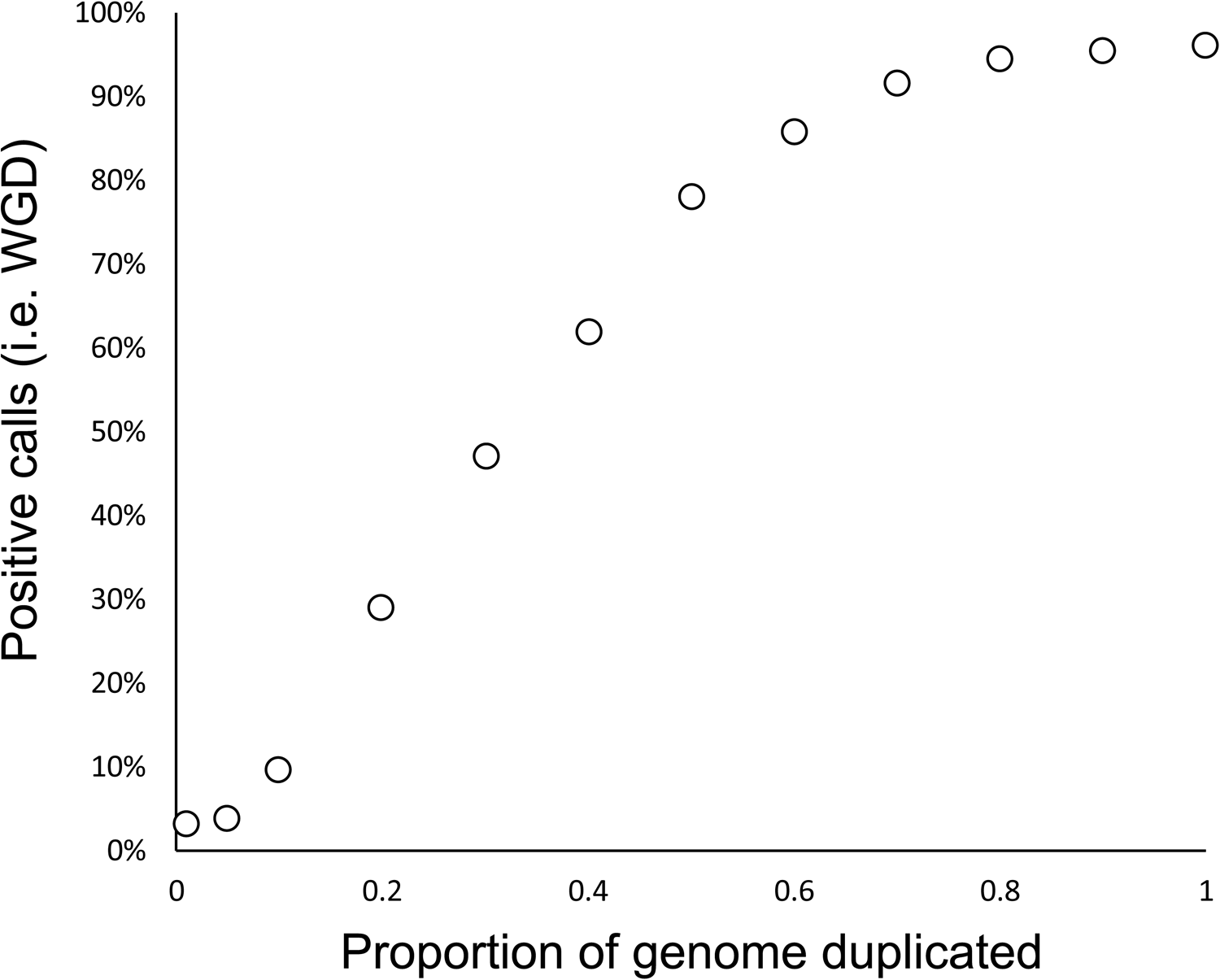
Effect of partial genome duplication on SLEDGe calls. Simulations modeled partial to full genome duplication ranging from 0.01 to 1 of the genome duplicated. Positive calls denotes the percentage of simulations at a given level of duplication that SLEDGe reported as depicting a WGD.

**Fig 5:**
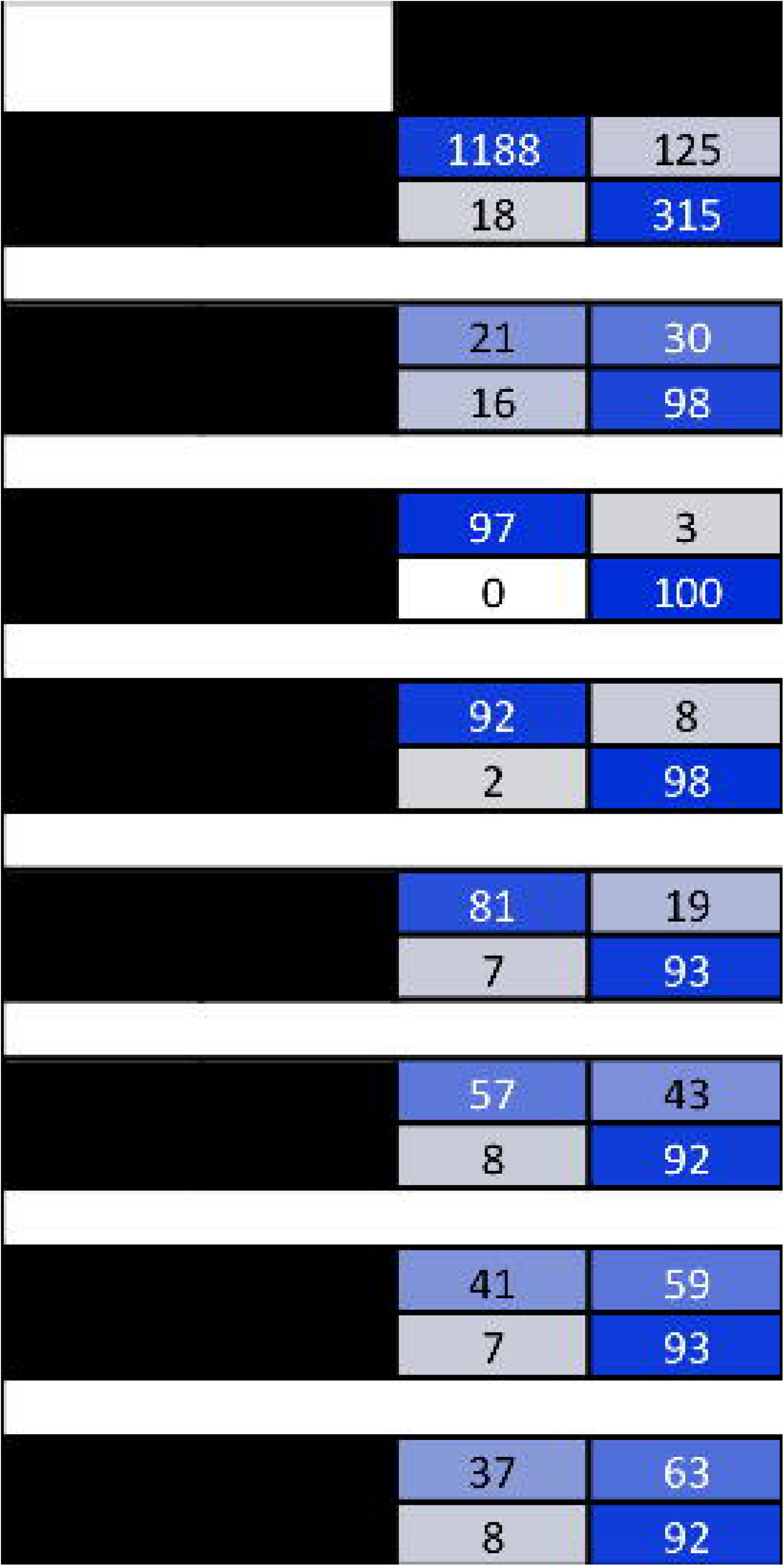
Effect of paralog number on SLEDGe calls. Both empirical samples from the 1KP dataset and simulated data were used.

We envision SLEDGe to be most useful for broad assessments of large numbers of taxa. In its current iteration, it can detect presence or absence of a WGD, but not estimate its age or other salient details, such as allo- vs autopolyploidy. It can, however, serve as a useful screening step before the implementation of more computationally costly methods, such as ksrates (Sensalari et al., 2022), MAPS (Li et al., 2015, Li et al., 2018), or WHALE (Zwaenepoel et al., 2019). This screening function can, however, be useful. We are careful in this paper to phrase SLEDGe’s performance on the full 1 KP dataset as “concordance” rather than accuracy. Most of the assessments of WGD in the 1KP paper were based on phylogenetic models and mixture models instead of synteny, and as such are also best considered estimates.

Areas where SLEDGe and these other techniques disagree are potentially interesting areas for further study. For example, SLEDGe finds no support for the CHROalpha duplication detected by other methods in the 1KP dataset. This duplication unites some *Chloromonas* algae species with *Chlamydomonas rienhardtii*. Extensive gene duplication has been found in this lineage that appear to be linked to stress-response genes (Wu et al., 2015). It is possible that unique circumstances that encourage large-scale but not whole-genome duplication may confound some WGD estimation methods. Likewise, SLEDGe fails to find WGD in the brown algae genus *Saccharina*, which are nested in a clade linked to the SAJAalpha duplication. As with Chloromonas, *Saccharina* has a history of large-scale tandem duplications (Ye et al., 2015), which may have confounded other estimation methods but perhaps not SLEDGe. By comparing these methods, points of disagreement may suggest fertile ground to investigate the evolution of large-scale but not whole-genome duplications.

We are continuing to improve the simulations upon which SLEDGe is based, including adding the ability to subset scaffolded genomic data to screen for aneuploidy and other large-scale duplications, determining allo- vs autopolyploidy, and identifying multiple duplication events. As phylogenetic and genomic methods and questions increase in both evolutionary depth and breadth, the need for methods to detect signals from large quantities of data only grows. The scope and effects of whole-genome duplication across plants, and more broadly all eukaryotes, is one of those questions. By incorporating SLEDGe as a tool for the rapid screening of ancient polyploidy in genomic and transcriptomic samples, especially as a first-pass before more computationally complex and time-consuming tools, we hope that investigators can spend less time painstakingly teasing out WGDs in their lineages of choice, and more time investigating the biological causes and implications of those duplications.

