## Supplementary figures and images for "SLEDGe: Inference of ancient whole genome duplications using machine learning"

### Supplemental Figure 1

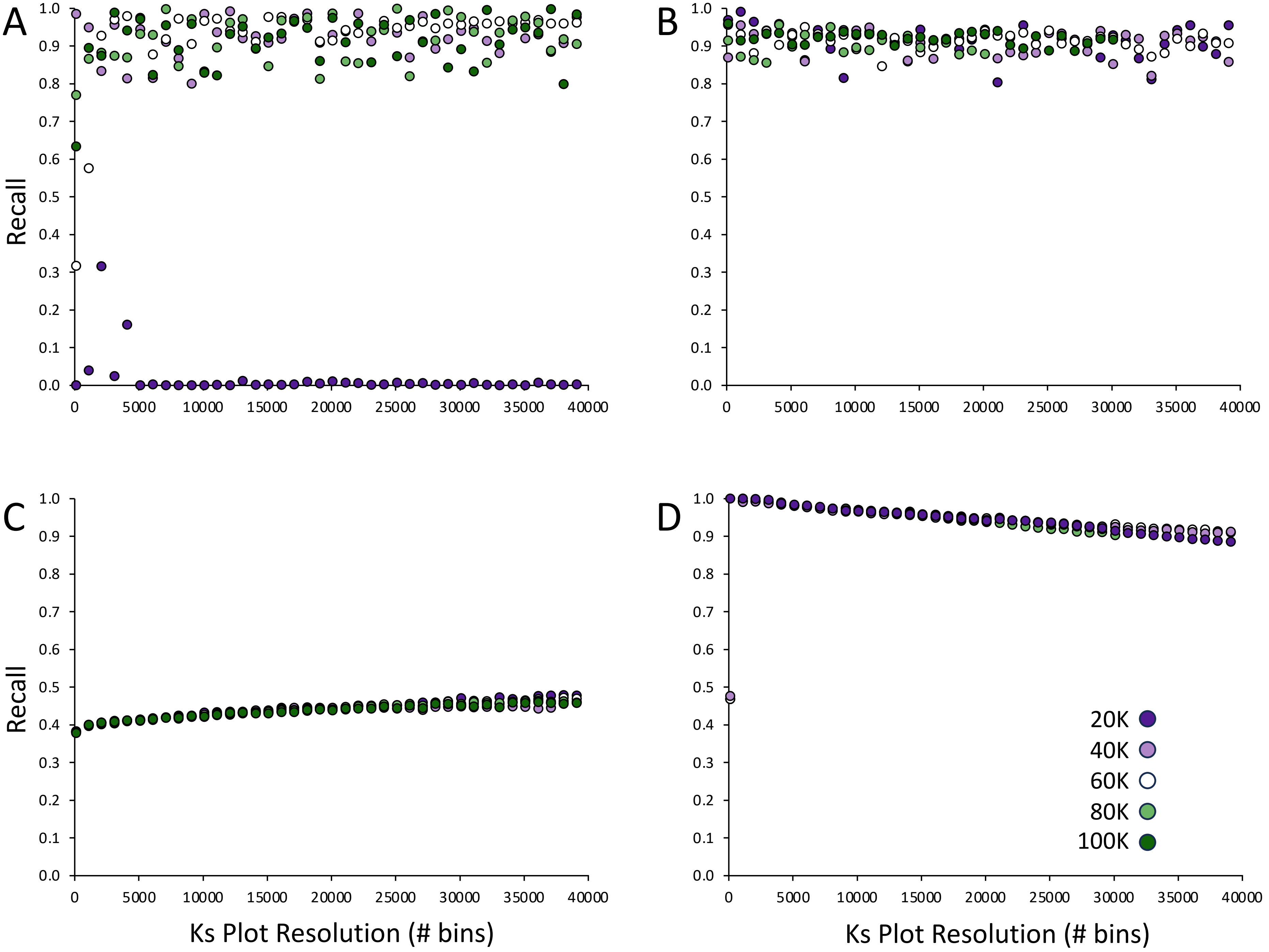

### Supplemental Figure 2

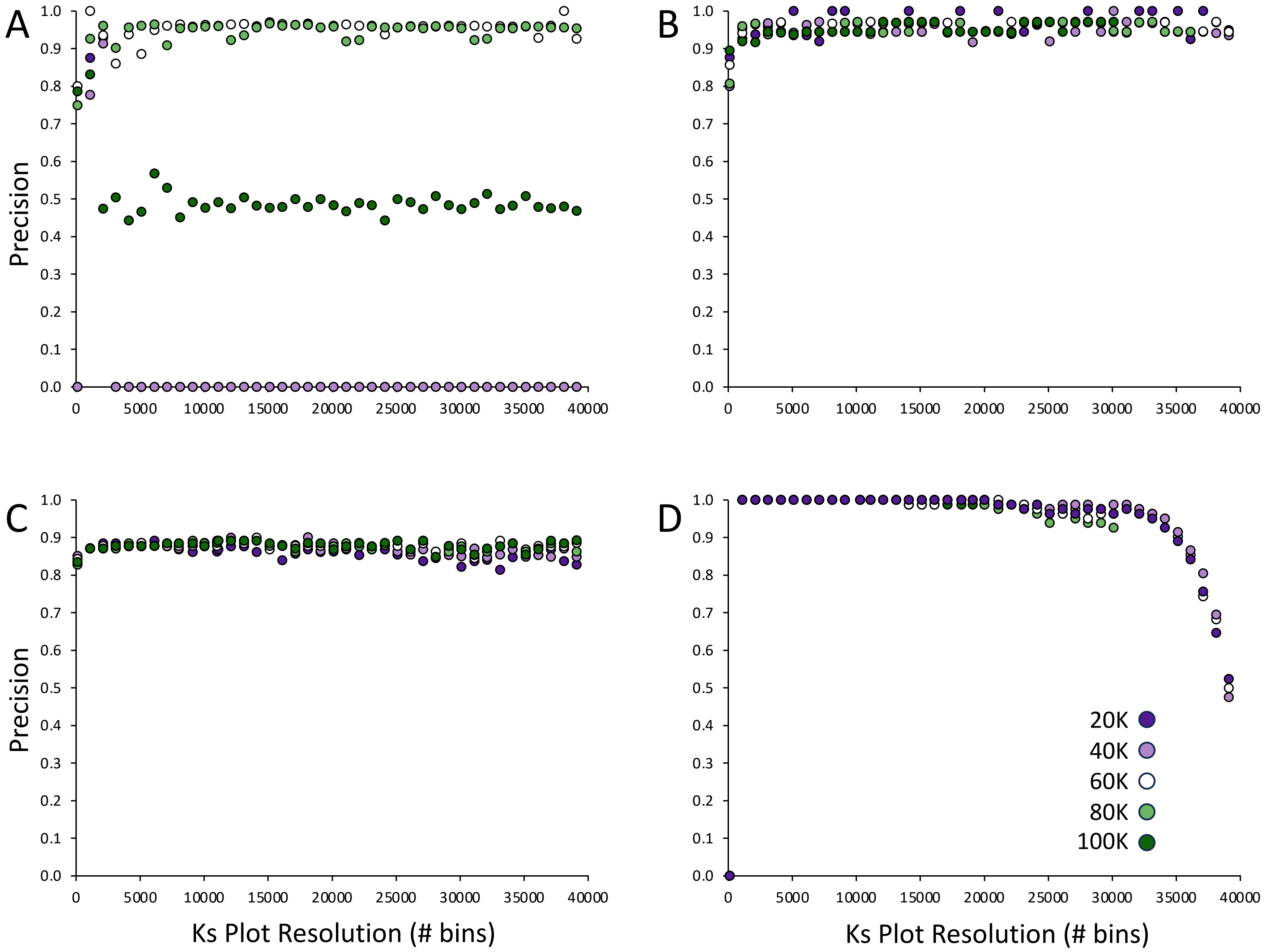

### Supplemental Figure 3

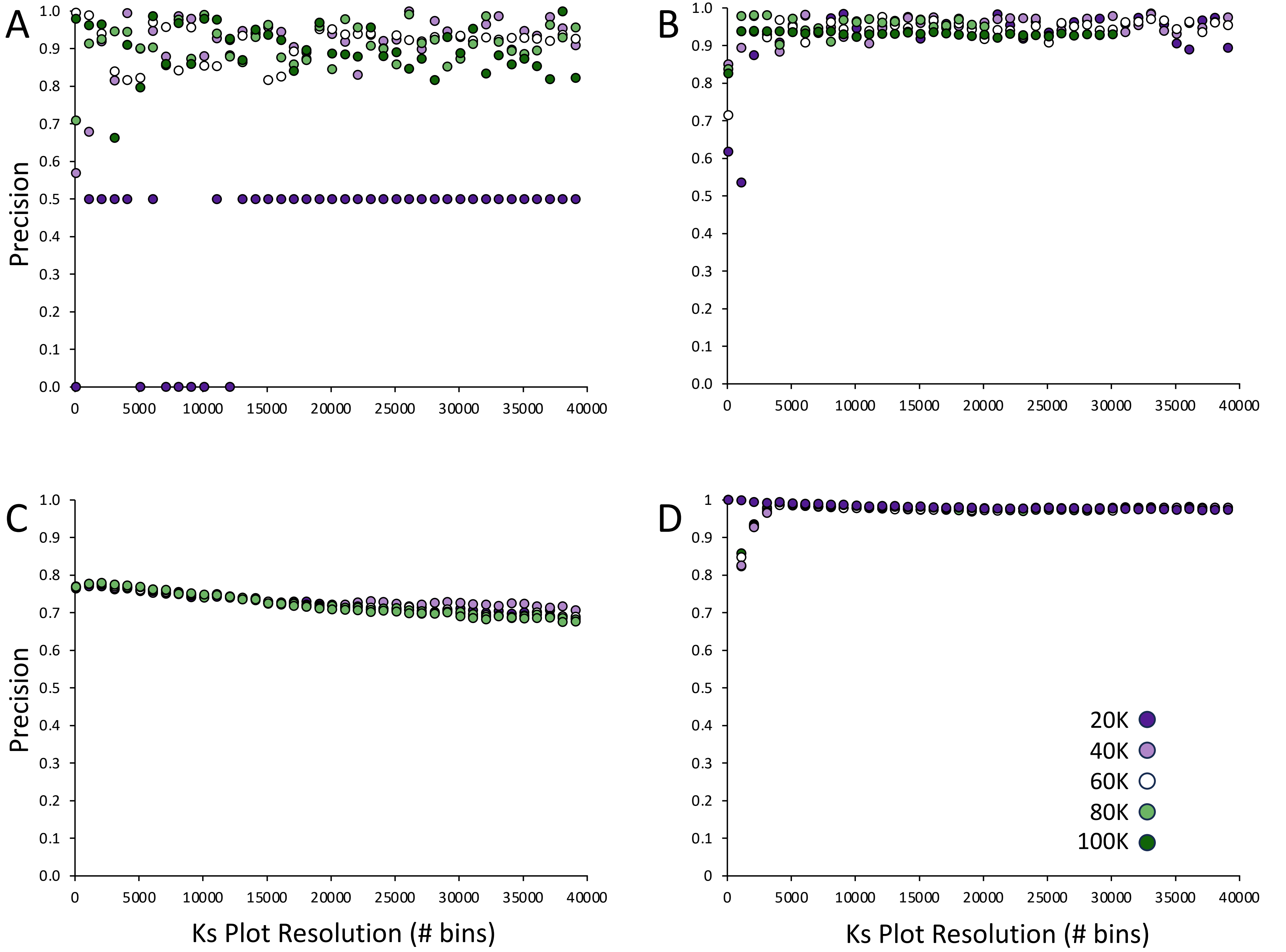

### Supplemental Figure 4

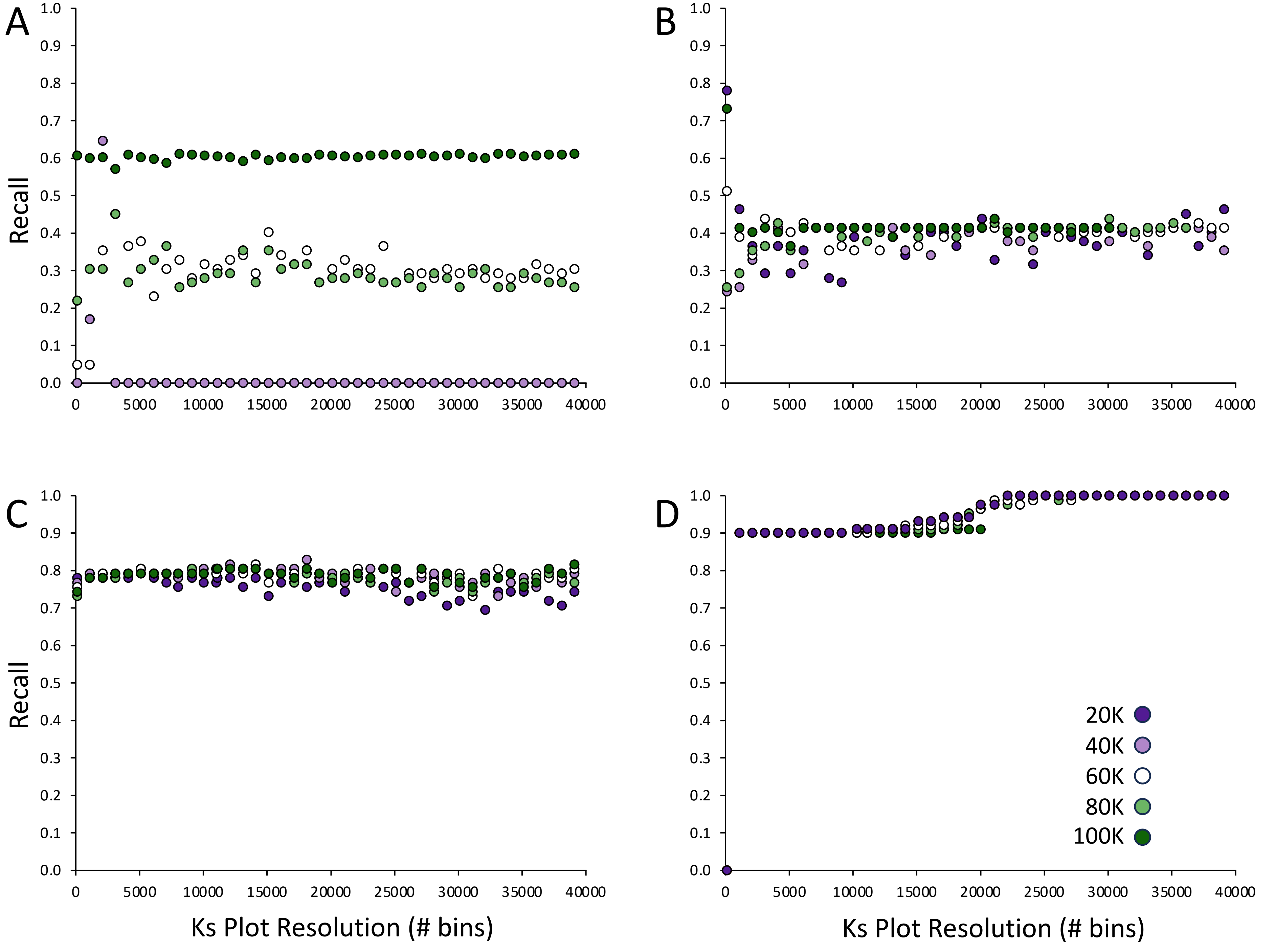
